# Farmed crickets raised with dermestids suffer from reduced and delayed growth, but not enough to explain reports of dramatic yield loss

**DOI:** 10.1101/2024.06.26.600772

**Authors:** MJ Muzzatti, MW Ritchie, EC Bess, SM Bertram, HA MacMillan

## Abstract

The mass production of crickets for food and feed is an expanding North American industry. Facilities that mass rear insects are at risk of pest infestations because the optimal environmental conditions for rearing beneficial species may also support the development of pest species. Here, we present the first recorded results detailing the interactions between dermestids and farmed crickets. Cricket farms have reported extremely low harvest yield during heavy dermestid infestations, but the exact reasons for this low yield are unknown. Many dermestid larvae are covered in dense, detachable, barbed setae called hastisetae, which are used by the larvae as an active trapping system against arthropod predators. We designed a series of experiments to test the hypotheses that dermestids (*Dermestes ater* DeGeer) may be directly impacting cricket (*Gryllodes sigillatus* Walker) yield through the physical effects of hastisetae ingestion and/or indirectly impacting cricket yield through competition for fishmeal, a primary source of protein in conventional cricket feed. Our predictions that cricket life history and survival would be negatively affected by dermestids were largely refuted. Females fed infested diets grew less mass, but not smaller body size, compared to females fed uninfested diets. We also found that while crickets experienced delayed growth early in life after living with dermestids, they were able to tolerate living with, and consuming, dermestid larvae. We discuss how these findings have led to new hypotheses concerning how dermestid infestations drive reductions in cricket farm yield.

## 1.0 Introduction

Mass rearing of insects for feed and food is still a developing industry, and while insects have been mass reared throughout history (e.g., honeybees, silkworms, parasitoid wasps), new species never previously reared *en mass* are now being farmed, such as mealworms, black soldier flies, and crickets (Larouche *et al*., 2023). This shift toward new species raises new challenges, one of the foremost being pest management (Deruytter *et al*., 2021). Mass reared insect facilities are at risk for the rapid proliferation of pest infestations because the environmental conditions optimized to rear the beneficial species, such as temperature, humidity, and feed, may also support the growth and development of other unwanted species (Deruytter *et al*., 2021). Managing insect pests in mass reared insect facilities presents a unique challenge, as pesticides are not a viable solution in an environment where the beneficial insects will undoubtedly be adversely affected, and the interactions between reared species and pest species are not well documented (Deruytter *et al*., 2021; Marshall, 2017). Very few pests of insect facilities for food and feed are recorded in the literature; pyralid moths (Pyralidae: Lepidoptera) infest mealworm (Tenebrionidae: Coleoptera) farms (Deruytter *et al*., 2021), and stored *Tenebrio molitor* (Tenebrionidae: Coleoptera) larvae meal is susceptible to infestation of stored-product beetle pests (Rumbos *et al*., 2020). To our knowledge, there is not any peer-reviewed published literature regarding pest interactions in cricket farm facilities.

Our personal communications with cricket farms have revealed that many cricket farms deal with beetle pests from the family Dermestidae. Most dermestid species are xerophilous necrophages – they develop on desiccated tissues and hairs of dead animals, and many are tolerant of hot, dry conditions with the ability to survive in substrate with very low water content (Marshall, 2018; Zhantiev, 2009). Cricket farms are an ideal environment for dermestids to thrive in, as they offer an unlimited source of animal protein from deceased crickets and shed exoskeletons, and often offer oviposition and pupation sites in wooden structures and the cardboard housing that is present in most farms. Extremely low harvest yields are reported by cricket farms during heavy dermestid infestations, but the exact reasons for low yields are unknown. Dermestids are known pests of stored grain facilities, homes, and museum collections, and one species, *Dermestes ater* DeGeer (Coleoptera: Dermestidae), is a recorded pest of silkworm (*Bombyx mori*) facilities in India (Geetha Bai and Mahadevappa, 1994; Kumar *et al*., 1990). *Dermestes ater* larvae bore into silkworm cocoons and feed on the pupae within (Ansari and Basalingappa, 1987; Kumar *et al*., 1988). Larvae will also bore and pupate through paper, cardboard, cork, wood and other construction materials (Roth and Willis, 1950; Zanetti *et al*., 2020). Adults of the same dermestid species feed on silkworm eggs and will oviposit directly into fishmeal and any available crevices (Kumar *et al*., 1990; Roth and Willis, 1950). Dermestids are also reared in captivity (for use in taxidermy) on diets consisting of mostly fishmeal (Coombs, 1981). Given that fishmeal is such a favorable feeding and egg laying substrate for dermestids, it is perhaps unsurprising that dermestids infest cricket farms, given fishmeal is commonly used as the main source of protein in cricket diets (Ssepuuya *et al*., 2021).

Many dermestid larvae are covered in dense, detachable, barbed setae called hastisetae, which are used by the larvae as an active trapping system against arthropod predators (Kokubu and Mills, 1980; Nutting and Spangler, 1969; Ruzzier *et al*., 2020). Dermestid hastisetae have been found obstructing the digestive tracts of invertebrate predators (Nutting and Spangler, 1969). Vertebrates, including humans, are also adversely affected by dermestids, with multiple reports of allergy, dermatitis, and lung and skin inflammatory reactions after dermestid exposure (Bernstein *et al*., 2009; Gumina and Yan, 2021; MacArthur *et al*., 2016; Simon *et al*., 2021). The internal structure of hastisetae contains an unidentified amorphous matrix, which is hypothesized to contain defensive proteins and/or chemicals, like the defensive hastisetae in Lepidoptera (Battisti *et al*., 2011; Ruzzier *et al*., 2021). However, dermestid hastisetae differ from Lepidoptera in that they do not have secretory or glandular cells (Ruzzier *et al*., 2022), and thus it is unlikely toxins are delivered through direct hastisetae contact. Hastisetae regenerate along with the epicuticle with each larval molt and the remaining shed moults contain the old hastisetae (Ruzzier *et al*., 2021). Mortality of weevils (Curclionidae) due to hastisetae-contaminated substrate has been recorded, suggesting hastisetae functionality persists over time even after they are shed during moulting and discarded (Ruzzier *et al*., 2021). It is unknown if dermestid hastisetae are harmful to crickets, and whether fully intact hastisetae or dermestid-infested cricket farm substrate (ex. feed, egg-laying medium) are responsible for reductions in cricket farming yield.

Understanding how pests interact with intentionally farmed species allows for the development of effective integrated pest management strategies that can target specific vulnerabilities in their life cycle. We hypothesized that dermestids may be directly impacting cricket (*Gryllodes sigillatus* Walker (Orthoptera: Gryllidae)) yield through the physical effects of hastisetae ingestion and/or indirectly impacting cricket yield through competition for fishmeal, a primary source of protein in conventional cricket feed. To test these non-mutually exclusive hypotheses, we designed two experiments to test 1) the effect of feeding crickets diets infested with hastisetae and whole dermestids, and 2) the effect of rearing both crickets and dermestids together in captivity with and without fishmeal. We measured and quantified survival of both species, food consumption, body size and mass at adulthood, and mass gain of *G. sigillatus*. We also sampled circulating hemocyte levels of *G. sigillatus* as a first step towards identifying whether crickets living with dermestids mount an immune response, which would be expected if crickets were being directly impacted by the hastisetae. Hemocytes play several roles in insect cellular immunity, such as phagocytosis, nodulation, encapsulation, and melanisation, but also serve many other roles in pathways unrelated to immunity (Stanley *et al*., 2023). We predicted that survival and life history traits would be severely reduced among crickets fed diets and dermestids with hastisetae, as well as crickets raised with dermestids and fishmeal. We also predicted that the removal of fishmeal from the diet would negatively impact dermestid survival and, as a result, would positively impact cricket performance and survival such that cricket response in the fishmeal-free treatments with dermestids would be comparable to the control with fishmeal and without dermestids. Finally, we predicted that crickets that were reared with dermestids would have higher circulating hemocyte levels compared to crickets that were not raised with dermestids.

## 2.0 Methods

### 2.1 Animal husbandry

All experimental crickets were received as eggs from a commercial cricket supplier (Entomo Farms, Norwood, Ontario, Canada), and a fresh batch of eggs was used for each experimental trial. The eggs were laid in a medium of peat moss and maintained inside an incubator (Thermo Fisher Scientific Inc., Massachusetts, United States) at 32°C and 60% relative humidity. A 14:10-hour light:dark cycle was maintained using an LED installed alongside the incubator interior. These conditions were maintained for all experiments. Live dermestids were received from local suppliers. Dermestids were identified as *Dermestes ater* using a pest identification guide and a dichotomous key (Biggs *et al*., 2022; Bousquet, 1990). Dermestids were either frozen immediately or were maintained separately from all laboratory-reared crickets at room temperature in a 22.9 x 15.2 x 15.2 cm (length, width, height) rearing container with egg cartons and *ad libitum* fishmeal. The walls of the container were misted with water every two days.

All crickets were photographed using a Zeiss Stemi 305 Stereo Microscope with an Axiocam 208 camera for body size measurements using ImageJ, version 1.48 software (Schneider *et al*., 2012). Body size measurements included head width, pronotum width, and pronotum length, and were averaged across three independent measurers to reduce measurement bias.

### 2.2 Hastisetae infested diets

Hastisetae were removed from dermestid larvae by first flash freezing 20 late-instar larvae inside a 15 mL Falcon tube with 10 g of cricket feed in liquid nitrogen for 10 s, and then vortexing the tube using a S|P ^®^ Vortex Mixer (cat. S8223-1, American Scientific Products, Michigan, United States) on setting = 10 for another 10 s. Hastisetae removal was confirmed through observation of bare larvae (Figure 4C). The resulting hastisetae infestation density of 3 larvae g^-1^ of feed was selected from reported feed board samples during heavy dermestid infestations at a cricket farm. Confirmation of hastisetae removal was performed through observation and removal of bare dermestid larvae from the tube. This process was repeated without dermestid larvae to create the control diet.

Individual three-week-old crickets were fed either diet *ad libitum* for five weeks. In the first trial, 40 crickets were fed each diet and in the second trial, 55 crickets were reared on the control diet and 45 on the infested diet. Survival and days to adulthood were monitored throughout the experiment, body size of each cricket was measured upon eclosion to adulthood, and mass was recorded at start of the experiment, at adulthood and 14 d post adulthood. This experiment was repeated twice, and during the second trial we measured weekly feed consumption. Principal component analysis was used to quantify adult body size by creating a single summary variable (PC1) for all three body size measurements. The first principal component explained 90.8% of the variation in size (eigenvalue = 2.73), and all size measures loaded equally on the first principal component. Statistical analyses for this experiment, including the principal component analysis, were performed using JMP^®^, Student Subscription 17.2.0 (SAS Institute Inc., Cary, NC, 1989-2023)

We performed a survival analysis to determine whether infested diets influenced survival. Individuals that died before the end of the experiment were treated as ‘failures’ and individuals that survived the experiment were treated as ‘censored’. We used linear models to determine whether infested diets influenced adult cricket body size, mass gain between the start and end of the experiment, and lifetime food consumption (total amount of food consumed/days survived). Diet, sex, and the interaction between diet and sex were included as fixed effects, and experiment trial was included as a random effect. Pairwise comparisons for the linear model were developed using Tukey’s Honestly Significant Difference test. We also used a linear mixed model to determine whether infested diets influenced cricket mass and amount of food consumed. Diet, sex, week, and all possible interactions were included as fixed effects, and experiment trial and cricket identification number were included as random effects. For amount of food consumed, only cricket identification number was included as a random effect because food consumption was exclusively measured in the second trial.

### 2.3 Whole dermestids feeding

To test whether crickets would consume entire dermestid larvae, and to determine whether they prefer dermestid larvae with or without hastisetae, we fed crickets freshly deceased late-instar dermestids with hastisetae (“Hairy”) or without hastisetae (“Hairless;” see hastisetae removal protocol above) to 30 four-week-old crickets for seven days. We chose this age because we wanted to isolate feeding effects from the chance of high mortality that smaller, early-instar cricket nymphs may experience when exposed to dermestid larvae. To control for the effect of feeding on dermestid larvae, we also reared 30 crickets on conventional cricket feed; these crickets did not receive any dermestids.

Dermestid larvae were frozen at -30°C upon delivery and each individual cricket was given two dermestid larvae on the first day of the experiment. Each cricket was checked daily, and if the cricket ate any part of one or both dermestid larvae, the larvae were replaced. Every two days, all dermestids were replaced to ensure fresh material was consistently provided. Survival and number of dermestids consumed was recorded each day, except for Day five which was missed due to human error. Crickets sometimes did not consume the entire dermestid larva; if there was approximately ≤ 0.5 a larva remaining, it was recorded as one larva consumed. If > 0.5 a larva was remaining, it was recorded as half a larva consumed. These measurements were approximated through observation and not precisely measured.

We ran a survival analysis to determine whether the diet treatments influenced survival using JMP^®^, Student Subscription 17.2.0 (SAS Institute Inc., Cary, NC, 1989-2023). The count data of dermestid larva consumption was not normal, and so we used the ‘MASS’ package in RStudio (version 2023.12.0, RStudio Team, 2020) to use a generalized linear mixed model with a quasi-Poisson distribution to determine if there was a difference in consumption over time between crickets fed hairy or hairless larvae. Diet and week were included as fixed effects, and cricket ID was included as a random effect. The interaction between diet and week was included in the original model, but it was not significant and was thus removed from the final model. A linear model was also fit to the total number of dermestids consumed per cricket fed each diet. Normality of total consumption data was confirmed through residual observation and through Shapiro-Wilk test (*P* = 0.53 Hairless; *P* = 0.22 Hairy).

### 2.4 Competition

To test the effect of rearing both species together in captivity with and without fishmeal, 30 crickets were reared in 22.9 x 15.2 x 15.2 cm (length, width, height) containers with or without 30 late instar dermestid larvae and with or without a diet containing fishmeal. A fifth treatment was included that contained 60 crickets and the fishmeal diet to control for the effect of insect density. Each treatment was replicated twice (two bins per treatment). Raw feed ingredients were obtained from a local feed mill (Campbellford Farm Supply Ltd., Campbellford, Ontario, Canada), and diets were mixed in laboratory using a stainless steel one-touch pulse control coffee grinder (Black & Decker, Maryland, United States) and a food processor (Cuisinart, Connecticut, United States). The control diet followed the same recipe as the conventional cricket feed, whereas the treatment diet did not include fishmeal and relative proportions of other ingredients were adjusted to ensure final protein:carbohydrate was relatively the same between the two diets (Supp Table S1). Late-instar dermestid larvae tended to pupate within a week of being placed into the humid incubators, and so to replicate a heavily infested farm environment, dermestid larvae were continuously replaced every week. At the end of each week, all remaining dermestids were removed from each container, their life stage and number of each life stage was recorded, and 30 fresh late-instar larvae were placed into each treatment container. Cricket and dermestid survival were recorded weekly for seven weeks, and bin weight was recorded weekly beginning at week three. After seven weeks, four individual crickets from each bin were randomly selected for hemocyte assays. Effort was made to select two male and two female, but in some bins < 4 crickets remained and the surviving ratio did not always reflect this.

Statistical analyses for this experiment were performed using JMP^®^, Student Subscription 17.2.0 (SAS Institute Inc., Cary, NC, 1989-2023). A linear mixed model was used to determine whether treatment affected bin weight and the proportion of crickets surviving. Treatment, week, and the interaction between treatment and week were included as fixed effects, and bin identification number and replicate were included as random effects. After running the mixed model on bin weight with all treatments included, it was clear that the N=60 crickets treatment was skewing the results; bin weight was always heavier in the N=60 crickets treatment compared to all other treatments because there was always more crickets. The mixed model for bin weight was then re-run without the N=60 crickets treatment. If an interaction term was significant, a customized F-test was run for each level of both factors in the interaction using ‘Test Slices’ in JMP^®^ with crickets reared without dermestids and with fishmeal as the treatment reference category.

### 2.5 Hemocyte assay

We adapted our hemocyte assay from a few different sources (Czarniewska *et al*., 2023; Drayton and Jennions, 2011; Hampton *et al*., 2021; Stoddart, 2011). Crickets were cold-anesthetized in Eppendorf tubes on ice. Fresh cricket saline solution was prepared (in mM: 150 NaCl, 5 CaCl_2_ Dihydrate, 2 NaHCO_3_, 40 glucose, 9 KCl, pH 7.2) and 44.8 mM citric acid was included to prevent coagulation of hemolymph. Thin wall glass capillary tubes (100ml long, 1.5 mm outer diameter, 1.12 mm inner diameter; Item #TW150-4) were marked 0.5075 cm along the length of the tube to indicate 5 μL of hemolymph to draw up. The junction where the abdomen connects to the thorax below the dorsal pronotum plate was pierced with a sterile 25-G needle parallel to the body to avoid piercing gut tissue, producing a clear bubble of hemolymph. Crickets were then gently squeezed to push out more hemolymph, and the microcapillary tube was used to draw up hemolymph, which was then pumped out and expelled into the dye and saline solution. A 1:3 hemolymph: buffer solution was used for mixing hemolymph, saline, and dye. For females and larger males, we used (5 μL of hemolymph:12.5 μL saline):17.5 μL dye, and for males, we used (2.5 μL hemolmyph:6 μL saline):8 μL dye). This is because females are larger than males, and 5 μL was consistently drawn from females and 2.5 μL from males. The solution was gently stirred with the microcapillary tube to ensure even distribution of the hemolymph. Next, 15 μL of solution was transferred into a hemocytometer well by micropipette, and a cover slip was placed on top. The hemocytometer was immediately photographed under a dissection microscope (Zeiss, Oberkochen, Stemi 508 with an axiocam 105 colour), capturing an entire 4×4 corner grid (Figure 5C).

Cell counting was performed using the Cell Counter plugin in ImageJ Fiji (Schindelin *et al*., 2012). Viable cells with intact membranes appeared as clear dots, and dead cells with damaged membranes that contained the trypan blue dye appeared as blue dots. Each photo was counted separately by two experimenters, and cell counts were averaged between the two. If counts differed by ≥10 cells, the photos were recounted. Cell concentration (cells/mL), live cell number, and viability were calculated using the following parameters in the Hemocytometer Calculator [https://www.hemocytometer.org/hemocytometer-calculator/]: number of squares = 1, initial square volume (mL) = 0.0001, big squares were counted, and dilution factor was adjusted based on 5 μL (dilution factor = 6.6) or 2.5 μL (dilution factor = 7) hemolymph samples.

A linear model was used in JMP^®^, Student Subscription 17.2.0 (SAS Institute Inc., Cary, NC, 1989-2023) to determine whether treatment, sex, or the interaction between sex and treatment affected hemocyte concentration. Replicate was included as a random effect. Tukey’s HSD was used for post hoc analysis of the treatment effect.

## 3.0 Results

### 3.1 Hastisetae infested diets

The probability of surviving to adulthood was unaffected by whether the diet was infested with hastisetae or not (Log-Rank *X*^2^ = 0.61, df = 1, *P* = 0.44; Figure 1); between the two diet trials, 96 total crickets died (Control = 54, Infested = 42) and 84 crickets survived to adulthood (Control = 41, Infested = 43). Time to adulthood was unaffected by diet treatment (*P* = 0.066; Table 1).

**Figure 1.**
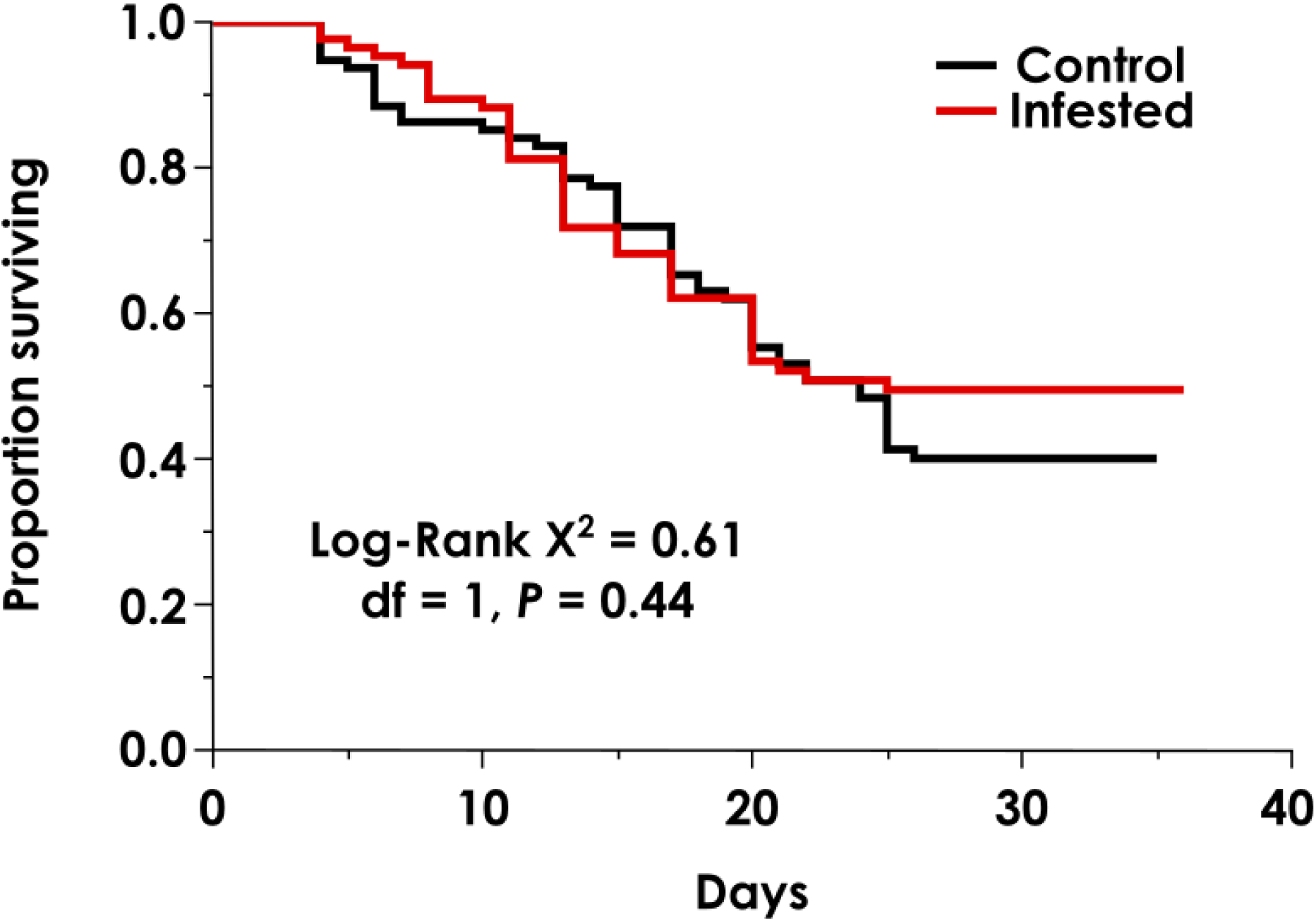
Kaplan-Meier plot representing the proportion surviving of N = 180 individual 3-week-old crickets fed one of two different diets: a control (0 dermestids/g diet) and dermestid hastisetae infested (3 dermestids/g diet) diet.

**Table 1.** Model results from linear models of how hastisetae infested diets influenced body size and mass gain of individual crickets. Terms in bold are statistically significant.

| Model | $R^2$ | Term | $df$ | $F$ | $P$ |
| --- | --- | --- | --- | --- | --- |
| Body size (PC1) | 0.61 | Diet | 1, 98.00 | 0.21 | 0.65 |
|  |  | **Sex** | **1, 98.02** | **141.64** | **<0.001** |
|  |  | Diet x sex | 1, 98.08 | 0.16 | 0.69 |
| Mass gain | 0.59 | Diet | 1, 54.01 | 3.35 | 0.073 |
|  |  | **Sex** | **1, 54.87** | **45.49** | **<0.001** |
|  |  | **Diet x sex** | **1, 54.01** | **5.23** | **0.039** |
| Lifetime food consumption | 0.40 | Diet | 1, 1.00 | 0.84 | 0.55 |
|  |  | **Sex** | **1, 1.00** | **35.55** | **<0.001** |
|  |  | Diet x sex | 1, 1.00 | 0.017 | 0.90 |
| Days to adulthood | 0.30 | Diet | 1, 101.00 | 0.069 | 0.066 |
|  |  | Sex | 1, 101.00 | 5.40 | 0.79 |
|  |  | Diet x sex | 1, 101.10 | 0.19 | 0.79 |

Hastisetae-infested diet did not affect adult cricket body size (*P* = 0.65), but females grew larger than males (*P* < 0.001) (Figure 2A; Table 1). The amount of mass gained between the start (mass at week 3) and the end of the experiment (mass at week 8) was significantly influenced by an interaction between sex and diet (*P* = 0.039; Figure 2B; Table 1): females gained 27.9% less mass during the experiment when they were fed the hastisetae-infested diet compared to when they were fed the un-infested diet; diet treatment did not, however, impact the amount of mass males gained during the experiment. The overall amount of food consumed was unaffected by diet treatment in both sexes (*P* = 0.55; Figure 2C; Table 2), and females ate 44.7% more food than males (*P* < 0.001).

**Figure 2.**
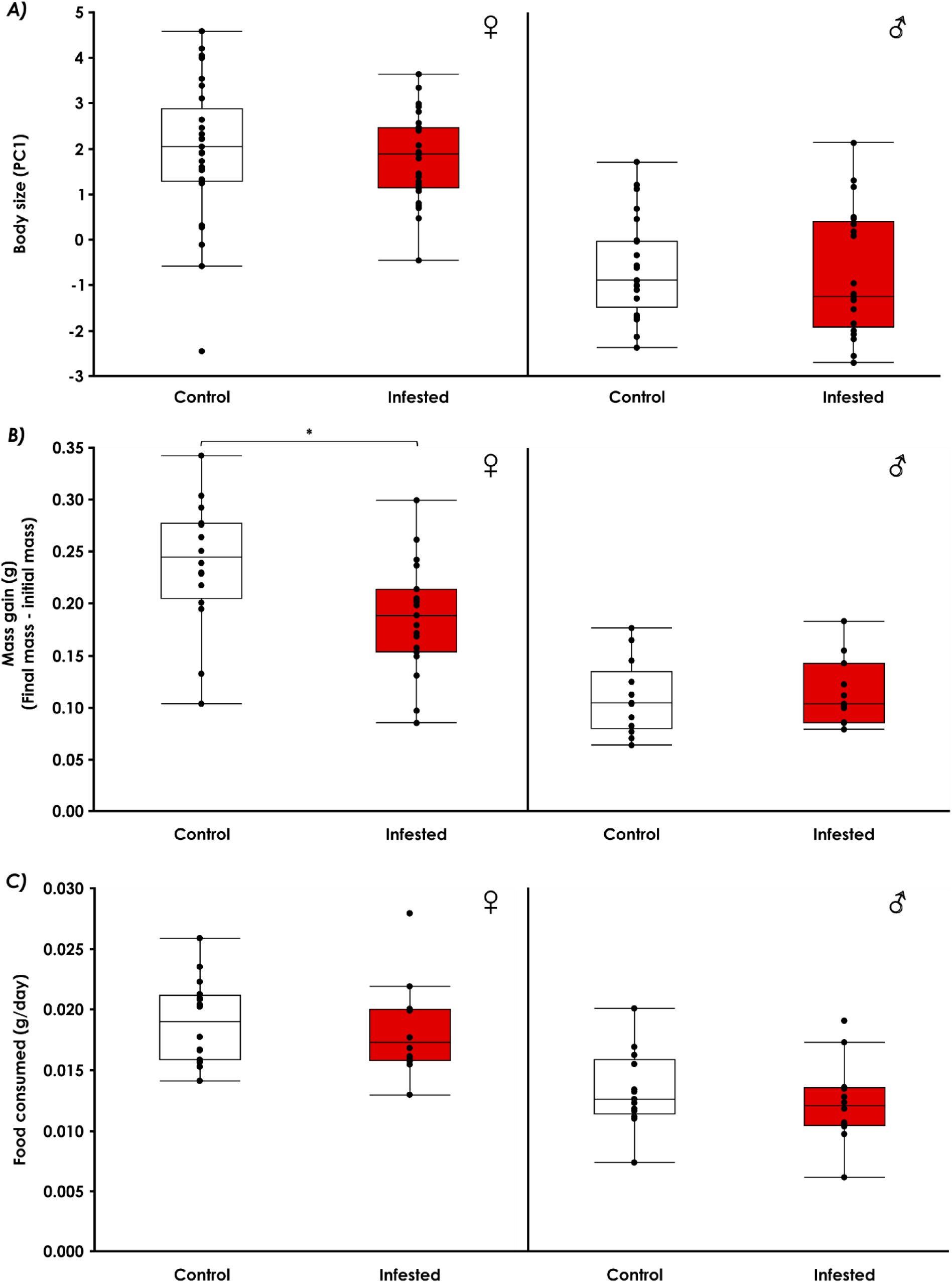
Adult life history traits of crickets fed one of two different diets: a control (0 dermestids/g diet) and dermestid hastisetae infested (3 dermestids/g diet) diet. Males and females were significantly different for each trait measured (*P* < 0.001). The centre line for each box represents the median, and the bottom and top lines represent the 25th and 75th quantiles. The whiskers extend 1.5 times the interquartile range. Solid circles represent individual data points. A) Body size (first principal component [PC1] built from head width, thorax width, and thorax length) of N = 84 adult crickets. B) Mass gain, calculated by subtracting the initial 3-week-old mass from the 8-week-old mass, of N = 59 adult crickets. C) The quantity of food consumed per day (total amount of food consumed divided by days survived) of N = 53 adult crickets.

**Table 2.**
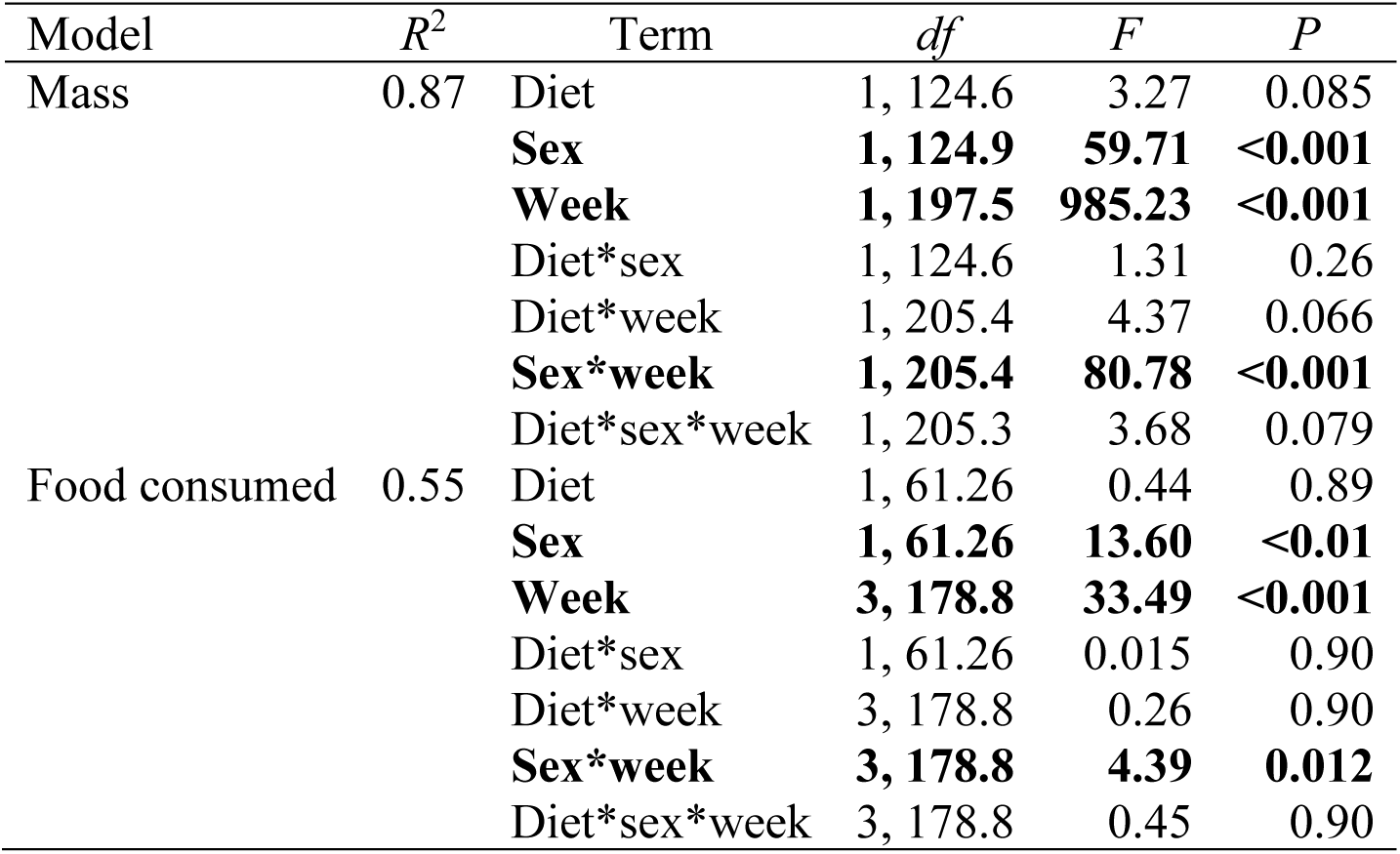
Summary statistics from repeated measures linear mixed models of mass and food consumption of individual crickets fed one of two different diets: a control (0 dermestids/g diet) and dermestid hastisetae infested diet (3 dermestids/g diet). Terms in bold are statistically significant.

| Model | $R^2$ | Term | $df$ | $F$ | $P$ |
| --- | --- | --- | --- | --- | --- |
| Mass | 0.87 | Diet | 1, 124.6 | 3.27 | 0.085 |
|  |  | **Sex** | **1, 124.9** | **59.71** | **<0.001** |
|  |  | **Week** | **1, 197.5** | **985.23** | **<0.001** |
|  |  | Diet*sex | 1, 124.6 | 1.31 | 0.26 |
|  |  | Diet*week | 1, 205.4 | 4.37 | 0.066 |
|  |  | **Sex\*week** | **1, 205.4** | **80.78** | **<0.001** |
|  |  | Diet*sex*week | 1, 205.3 | 3.68 | 0.079 |
| Food consumed | 0.55 | Diet | 1, 61.26 | 0.44 | 0.89 |
|  |  | **Sex** | **1, 61.26** | **13.60** | **<0.01** |
|  |  | **Week** | **3, 178.8** | **33.49** | **<0.001** |
|  |  | Diet*sex | 1, 61.26 | 0.015 | 0.90 |
|  |  | Diet*week | 3, 178.8 | 0.26 | 0.90 |
|  |  | **Sex\*week** | **3, 178.8** | **4.39** | **0.012** |
|  |  | Diet*sex*week | 3, 178.8 | 0.45 | 0.90 |

In the mixed models that examined how individual mass and food consumption changed over time, diet treatment did not significantly influence mass (*P* = 0.085; Figure 3A) or amount of food consumed (*P* = 0.70; Figure 3B; Table 2). Females ate more food than males (*P* < 0.0001) and the overall amount of food consumed changed over time (*P* < 0.0001; Figure 3B; Table 2). Somewhat unexpectedly, the three-way interaction between diet, sex, and week did not significantly influence mass (P = 0.079; Table 2). Trial was never a significant source of variation (*P* > 0.05), and individual differences among crickets (cricket ID) was always a significant source of variation (*P* < 0.05).

**Figure 3.**
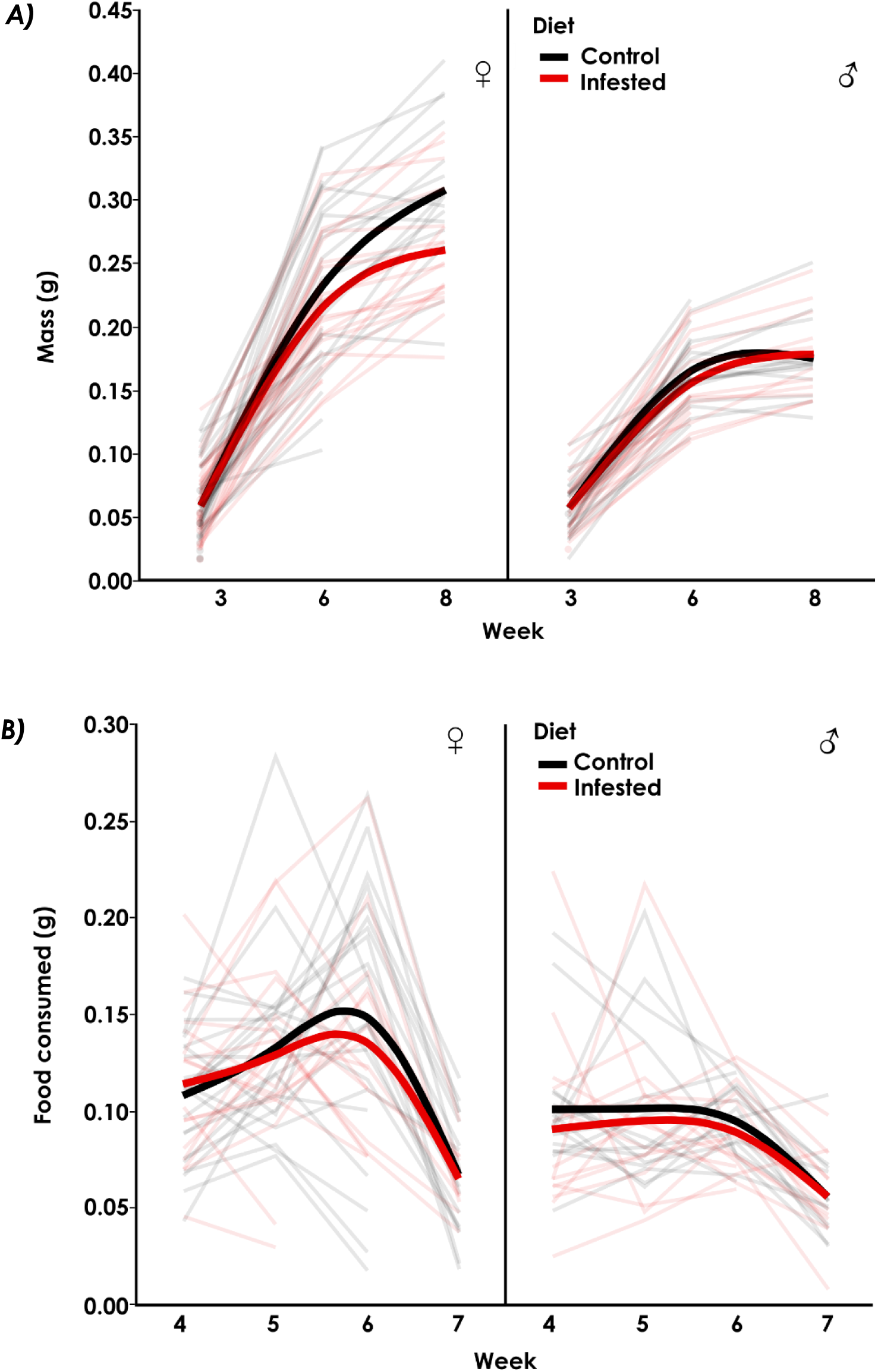
Development of mass as a function of age (A) and weekly food consumption (B) of individual crickets fed one of two different diets: a control (0 dermestids/g diet; N = 95) and dermestid hastisetae infested diet (3 dermestids/g diet; N = 85). Males and females were significantly different (*P* < 0.001). Smoothed thick lines connect mean values at each time point, and individual crickets are represented by thin lines over time. Solid dots in (A) represent individual crickets that did not survive past week 3.

### 3.2 Whole dermestid larvae

The probability of cricket survival was unaffected by feeding on whole dermestid larvae (Log-Rank *X*^2^ = 2.49, df = 1, *P* = 0.290); 9 crickets died (Control = 5, With hastisetae = 2, Without hastisetae = 2), and a single cricket on the control diet went missing. There was a significant effect of diet in the mixed model that evaluated factors influencing daily number of dermestids consumed: crickets that were fed hairy dermestids ate less than crickets fed hairless dermestids (*P* = 0.021; Figure 4A; Table 3). In total, crickets fed hairy dermestids ate ∼23% less compared to crickets fed hairless dermestids (F_1,58_ = 4.32; *P* = 0.042; Figure 4B; Supp. Table S2).

**Figure 4.**
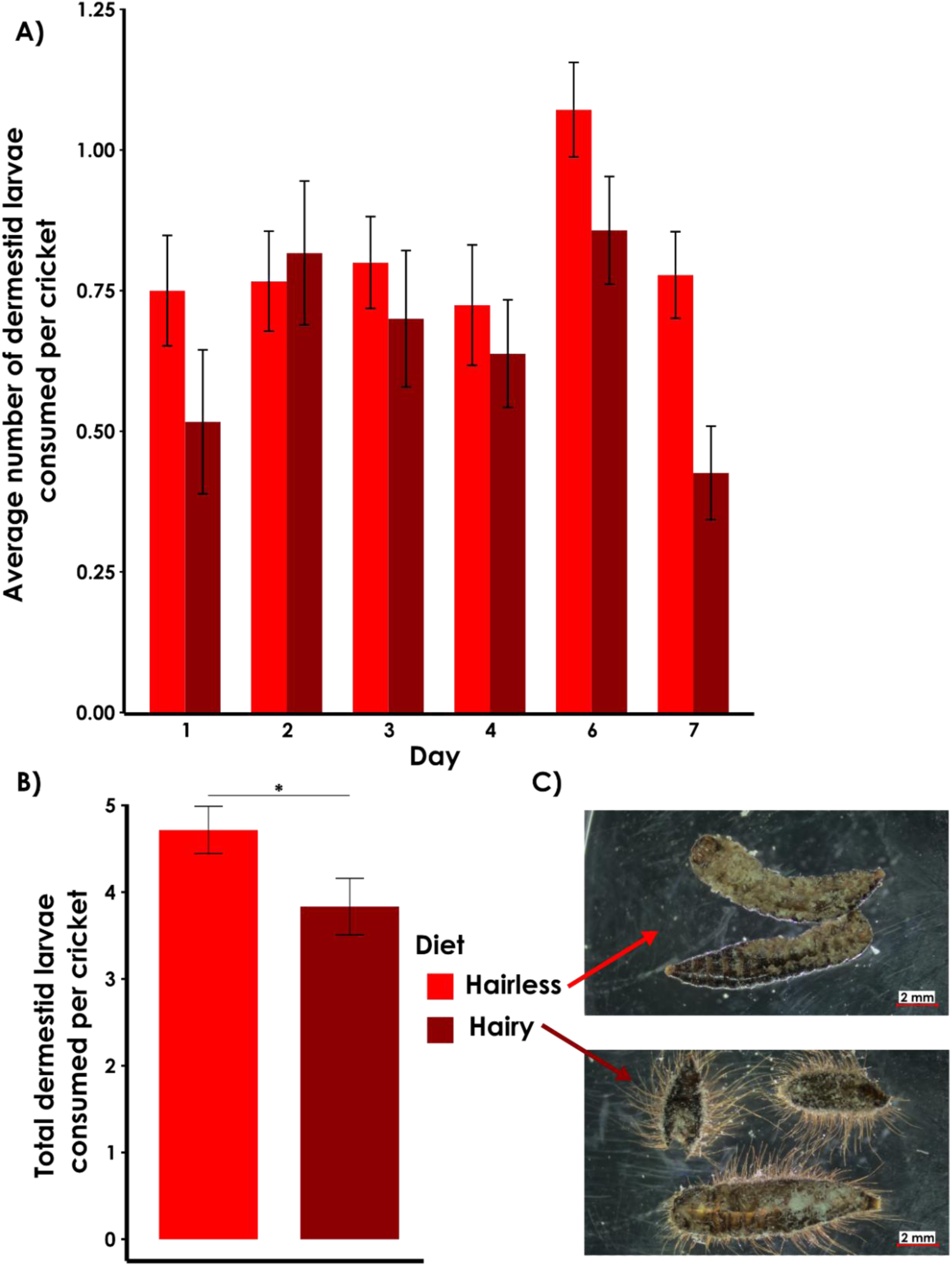
A) Average number of late-instar dermestid larvae consumed per cricket per day. Error bars represent standard error of the mean. Day 5 was missed due to human error, and thus not included on the figure. B) Total dermestid larvae consumed by N = 30 crickets on each diet treatment. C) Hairless dermestids represent dermestid larvae with their hastisetae removed, whereas hairy dermestids had their hastisetae unaltered. Larvae in photos were recovered after cricket feeding.

**Table 3.**
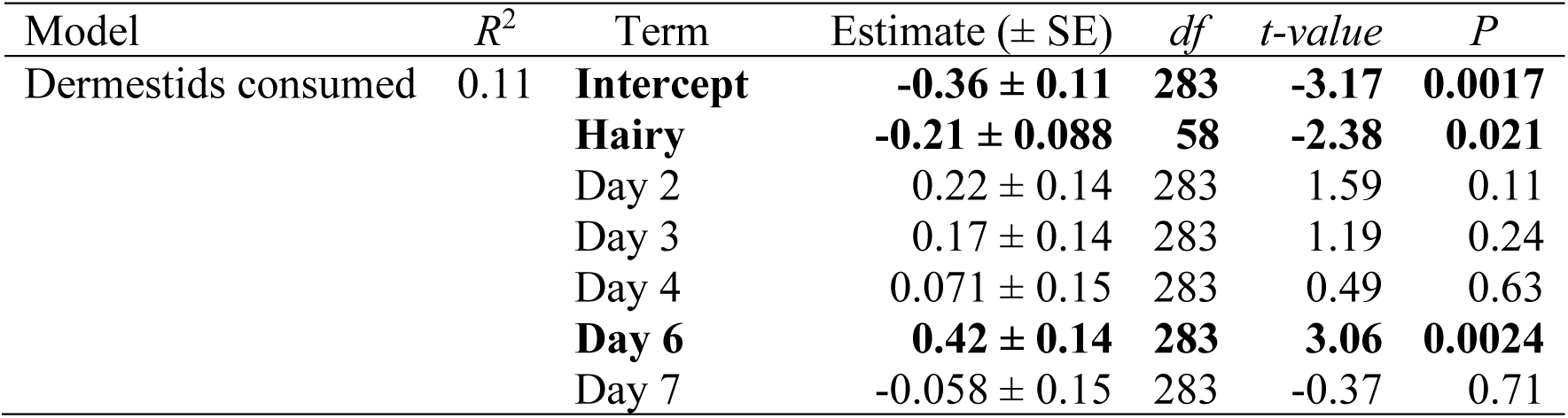
Summary statistics from the generalized linear mixed model of food consumption of individual crickets fed hairy and hairless dermestids for seven days. Hairless dermestids represent dermestid larvae with their hastisetae removed, whereas hairy dermestids had their hastisetae unaltered. The *R*^2^ represents the conditional *R^2^* for lognormal residual distribution. Day 5 was missed due to human error, and thus not included in the analysis. The intercept represents Diet = Hairless and Day = 1. Terms in bold are statistically significant.

| Model | $R^2$ | Term | Estimate (± SE) | $df$ | $t$-value | $P$ |
| --- | --- | --- | --- | --- | --- | --- |
| Dermestids consumed | 0.11 | **Intercept** | **-0.36 ± 0.11** | **283** | **-3.17** | **0.0017** |
|  |  | **Hairy** | **-0.21 ± 0.088** | **58** | **-2.38** | **0.021** |
| | | Day 2 | 0.22 ± 0.14 | 283 | 1.59 | 0.11 |
| | | Day 3 | 0.17 ± 0.14 | 283 | 1.19 | 0.24 |
| | | Day 4 | 0.071 ± 0.15 | 283 | 0.49 | 0.63 |
|  |  | **Day 6** | **0.42 ± 0.14** | **283** | **3.06** | **0.0024** |
| | | Day 7 | -0.058 ± 0.15 | 283 | -0.37 | 0.71 |

### 3.3 Competition

The proportion of crickets surviving significantly decreased over time in the competition experiment (*P* < 0.001), however there were no significant effects of treatment or any interactive effect of week and treatment on survival (Figure 5A; Table 4). There was also no overall effect of treatment on bin weight (*P* = 0.97; Table 4), but there was a significant interaction between treatment and week on bin weight (*P =* 0.023; Table 4). At week three, crickets reared with just fishmeal had ∼92% significantly heavier bin weight compared to crickets reared with both dermestids and fishmeal (t = -2.83; *P* = 0.031; Figure 5B).

**Figure 5.**
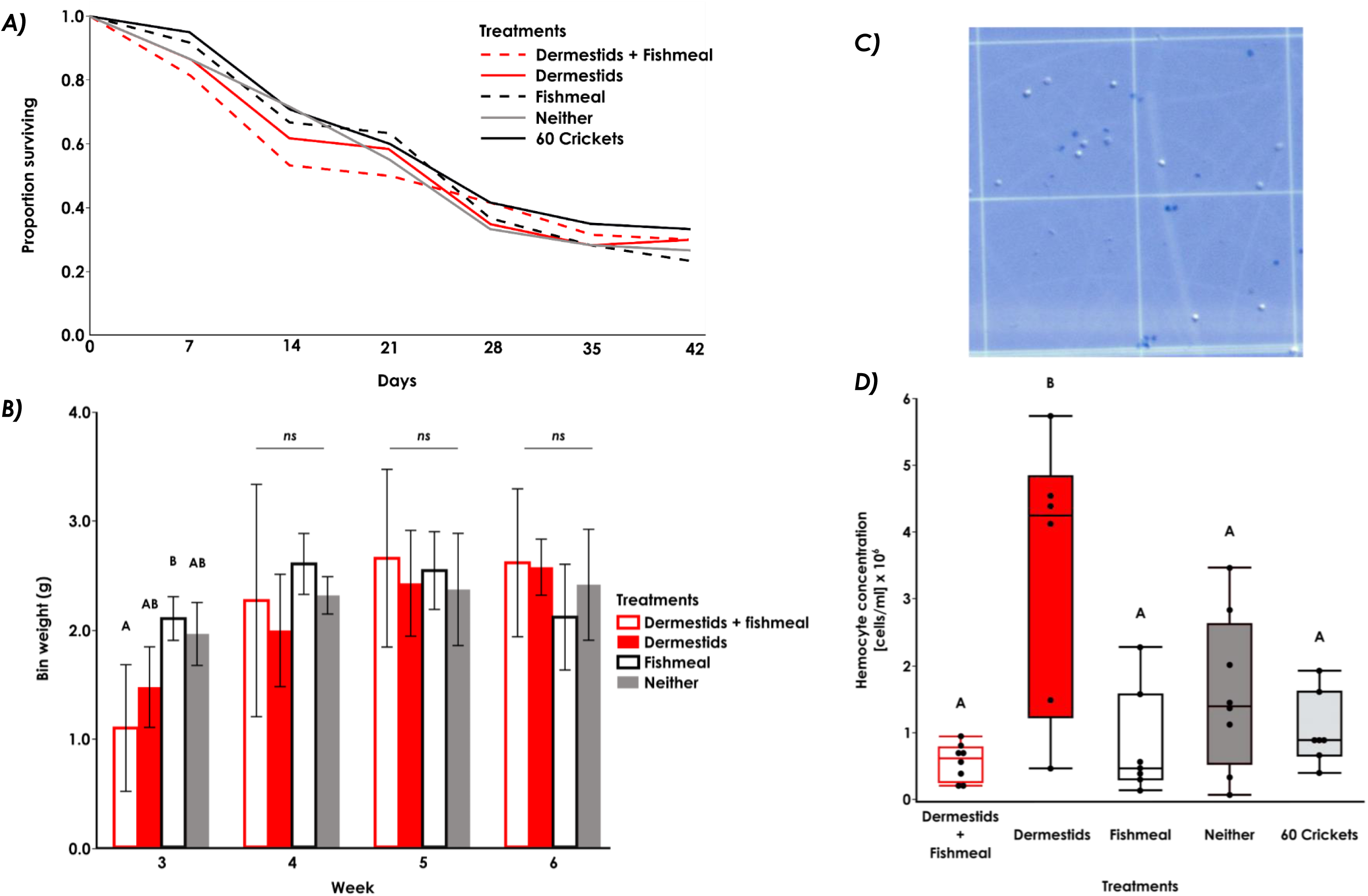
A) Kaplan-Meier plot representing the proportion of N = 30 crickets after 6-weeks living with or without late-instar dermestid larvae and fed a diet with or without fishmeal. B) Bin weight (total cumulative weight of crickets in each bin; N = 2 bins per treatment) of crickets living with or without late-instar dermestid larvae and fed a diet with or without fishmeal. Bars represent standard error, and bars that do not share a letter within a week significantly differ, and *ns* represents no significant differences. C) A hemocytometer photograph of viable and dead cells from the competition experiment of crickets living with and without live dermestids and a diet containing fishmeal. Viable cells with intact membranes appeared as clear dots, and dead cells with damaged membranes that contained the trypan blue dye appeared as blue dots. D) Circulating hemocyte concentration [cells/mL] of crickets living for six weeks with or without late-instar dermestid larvae and fed a diet with or without fishmeal. The centre line for each box represents the median, and the bottom and top lines represent the 25^th^ and 75^th^ quantiles. The whiskers extend 1.5 times the interquartile range. Solid circles represent individual data points, and an auto jitter was applied along the x-axis to visually separate similar values. The 60 crickets control was included to control for density of insects within a bin, and included fishmeal but no dermestids.

**Table 4.** Summary statistics from the linear mixed models of bin weight and proportion surviving of crickets living with or without late-instar dermestid larvae and fed a diet with or without fishmeal. The 60 crickets control was included to control for density of insects within a bin, and included fishmeal but no dermestids. The bin weight model did not include the N=60 crickets control, but the proportion surviving model did. Terms in bold are statistically significant.

| Model | $R^2$ | Term | $df$ | $F$ | $P$ |
| --- | --- | --- | --- | --- | --- |
| Bin weight | 0.92 | Treatment | 3, 3 | 0.24 | 0.87 |
|  |  | **Week** | **3, 12** | **23.17** | **<0.001** |
|  |  | **Treatment\*week** | **9, 12** | **3.95** | **0.023** |
| Proportion surviving | 0.95 | Treatment | 3, 3 | 0.077 | 0.97 |
|  |  | **Week** | **6, 24** | **149.50** | **<0.001** |
|  |  | Treatment*week | 18, 24 | 0.96 | 0.79 |

Crickets reared with dermestids and without fishmeal had a significantly higher circulating hemocyte concentration compared to all other treatments (*F*_1,25.20_ = 10.81; *P* < 0.001; Figure 5D; Supp. Table S3). Female crickets also had a higher hemocyte concentration compared to males (*F*_4,25.60_ = 6.57; *P* = 0.025; Supp. Table S3), but there was no significant interaction between treatment and sex (*F*_4,25.22_ = 1.78; *P* = 0.16; Supp. Table S3).

## 4.0 Discussion

This series of experiments is the first to directly examine potential direct interactions between dermestids and mass reared crickets. These experiments were motivated by reports of severe reductions in cricket farm yield under heavy dermestid infestations, so we predicted severe reductions in survival, mass, and body size of crickets fed and/or reared with dermestid larvae. However, our predictions were refuted, as our effects were mild not severe. Crickets interacted with and fed on dead dermestids when given no other choice of food, and overall, survival and life history responses were mostly unaffected by feeding on infested feed and whole dermestid larvae. The only clear negative impacts of dermestids on crickets we observed here were a sex-specific reduction in mass gain of females fed diets infested with hastisetae (Figure 2B), a reduction in bin weight in early-life for crickets living with dermestids and fed a diet that included fishmeal (Figure 5B), and an increase in hemocyte concentration (indicative of an immune response) in crickets living with dermestids, but only while being fed a diet that included fishmeal (Figure 5D). Female crickets fed hastisetae infested diets were ∼28% lighter, but not smaller in size, compared to female crickets fed uninfested diets (Figure 2A,B). We have reported a similar result in this species; after feeding *G. sigillatus* a royal jelly dietary supplement, females grew heavier mass but not larger body size, a result explained by significantly more eggs found in royal jelly-fed females (Muzzatti *et al*., 2022). Dermestid infestations may reduce the number of eggs female crickets produce and oviposit, which could severely restrict the number of offspring contributing to subsequent generations and result in dramatic yield loss. We were unable to test this hypothesis and measure egg production in our crickets because the samples were used for unsuccessful attempts to filter and quantify hastisetae from cricket gut tissue. Future experiments should measure how egg production is influenced by dermestids. Regardless, females also ate more food than males in the infested diets experiment, thus ingesting more hastisetae, and so this may have been the driving factor behind the mass gain reduction. If we had tested a higher infestation density, we may have also observed a similar effect in males.

Our individual-level experiments feeding crickets infested diets and whole dermestid larvae were intentionally started using three- and four-week-old crickets to avoid high rates of mortality unrelated to dermestids if we started with younger, fragile cricket nymphs. Crickets at intermediate ages appear to tolerate the hastisetae density of three dermestids per gram of feed, as well as being provided with two dermestid larvae a day for food. This suggests that crickets can tolerate ingesting dermestids, regardless of whether dermestid hastisetae are in the food or dermestids are the only available food. However, there appeared to be a delay in growth among crickets living with dermestids and fishmeal in the diet. By the end of the experiment, crickets had largely recovered and there were no strong differences in bin weight among the treatments. Testing a higher density of dermestids may have resulted in a stronger effect, however this experiment required many fresh, late-instar dermestid larvae weekly, and we used the maximum number available to us. We tried to replicate a constant presence of dermestids throughout the experiment by adding new dermestids every week, but under a severe infestation in a farm-setting, especially one that uses wooden structures, there could exist overlapping generations of diapausing dermestids resulting in a constant pressure throughout successive generations of crickets. Our results hint at early-life reduction in mass of crickets living with dermestids, and so future research on interactions between these species could focus on how crickets in the vulnerable first few instars respond to living with dermestids.

From the whole dermestid feeding experiment, we learned that although crickets will readily consume entire dermestid larvae when given no other food options, they demonstrate a feeding aversion to the hastisetae as they consumed ∼23% less larvae that still had their hastisetae attached compared to the ‘hairless’ dermestid larvae (with their hastisetae removed; Figure 5.7B). This finding could indicate that cricket feeding on dermestid larvae with intact hastisetae is risky, and that crickets in an infested-farm environment are likely to avoid dermestid larvae and spend less time feeding in dermestid-infested feed. The primary function of hastisetae is an active trapping mechanism facilitated by density of setae and by the sharp pointed tip and barbed shaft (Supp. Figure S1; Kokubu and Mills, 1980; Nutting and Spangler, 1969; Ruzzier *et al*., 2020). We observed specific feeding damage between the hairy and hairless treatments; among the fed-upon hairy dermestids, feeding damage was observed along the ventral surface of the larvae which is sparser in hastisetae compared to the dorsal surface (Figure 5.7C). Often, the posterior halves of the hairy larvae were left behind, hollowed-out from a small feeding hole but hastisetae mostly left intact (Figure 5.7C), whereas entire hairless larvae were often completely consumed. An interesting follow-up experiment would be to feed crickets dermestid larvae starting earlier in life (when crickets are smaller and thus may not be as adept at dealing with impaling hastisetae) and using scanning electron microscopy to photograph the mouthparts of crickets to detect if hastisetae impale or impair mouthparts or other areas of the body. Regardless, it is clear that crickets consume dermestid larvae, a dangerous feeding activity for other invertebrate predators (Nutting and Spangler, 1969) and one that may also be dangerous for crickets past a certain quantity of larvae consumed.

A disposition by crickets to consume dermestid larvae may explain why the total combined mass of crickets in the bins (bin weight) living with dermestids but without fishmeal was not significantly different from the other treatment combinations. Assuming that dermestids were negatively affected by the lack of fishmeal, crickets may have fed upon struggling or deceased dermestid larvae. The number of dermestids recovered weekly from each bin decreased over time and fewer dermestids were recovered from bins without fishmeal (Supp Figure S3) but it is unclear whether crickets were consuming live dermestids, if dermestids were cannibalising each other, or dermestids were dying and scavenged by both crickets and dermestids.

The significantly higher hemocyte concentration in crickets that were living with dermestids but without fishmeal (Figure 5.10B) lends partial support to the idea that in the absence of fishmeal crickets were consuming more dermestids, but only if consuming large numbers of dermestids is stressful and elicits an immune response. Hemocytes have many roles in insect cellular immunity but also in non-immunity pathways (Stanley *et al*., 2023). Therefore, insect immune function is best measured using an integrative approach; for example, combining results of antibacterial activity, hemocyte microaggregations, and circulating hemocytes (Duffield *et al*., 2018, 2019) or measuring active phenoloxidase activity which plays a major role in humoral and cellular immunity and helps with wound closure and repair (Stanley *et al*., 2023). Due to time constraints, we were unable to follow up with other immunity-measuring approaches, but an experiment that combines all these approaches would be valuable to test whether dermestids do elicit an immune response of crickets, and whether this response is diet-dependent.

Behavioural experiments testing the trapping mechanism of dermestid hastisetae on crickets would be an interesting next step towards understanding the interactions between dermestids and crickets that drive reductions in production yield. Dermestid larvae feed and tunnel through cricket feed, and there is a possibility that crickets consume less dermestid-infested feed and/or spend less time feeding when the feed is dermestid infested. It remains unclear if, or how often, crickets are trapped by dermestid hastisetae under higher densities. In our experiments, only a single observation was recorded of an adult male cricket that became entangled in dermestid hastisetae and died (Supp. Figure S3). This observation is in stark comparison to an experimental accident reported by Ruzzier et al. (2021). Their dermestid rearing substrate was infested by weevils (*Sitophilus* spp.; Curculionidae), a beetle family notorious for extremely thick outer cuticle (Anbutsu *et al*., 2017). All the weevils were discovered deceased although the weevils never encountered live larvae, just old rearing substrate (Ruzzier *et al*., 2021). The structure of dermestid hastisetae varies among species (Supp. Figure S1; Ruzzier *et al*., 2021), and so some species may be more adept at entangling predators than others. Regardless, avoidance behaviour by crickets may provide a simple explanation to the reduction in yield of cricket farms under heavy dermestid infestations. Crickets may be avoiding feed trays infested with dermestids, consuming less food, and ultimately not growing as fast or large than if they were feeding regularly.

The dynamic nature of any new business or industry can often outpace the science required to solve complex problems as they arise. Dermestid infestations in cricket farms are a good example of such a challenge. Therefore, a discussion of existing and potential avenues of control methods is warranted to streamline the development of an integrated pest management program. Control of dermestid beetles in insect production facilities is difficult to achieve, and consists mainly of maintaining hygiene, performing heat treatments, and applying fumigants and contact insecticides (Kumar *et al*., 1990; Wilches *et al*., 2016). A variety of insecticides are suggested for chemical control (Wilches *et al*., 2016), but insecticides are not a desirable management solution when they may also impact the farmed insect. Larvae can cause structural damage and remain hidden from detection and insecticide treatments by tunneling into wooden structures (Zanetti *et al*., 2020). It is therefore strongly recommended to avoid wooden structures within cricket rearing facilities to prevent the spread of dermestid infestations. Clever application of materials science can help mitigate the influence of pests in farms; larvae of *D. maculatus* are unable to climb plastic surfaces (Geden and Carlson, 2001), but maintaining hygiene is crucial to prevent the build-up of dirt and particulate matter that may assist in climbing behaviour. Many dermestid species can enter diapause as larvae which can synchronize development of individuals within a population and can increase survival during extreme temperatures (Osuji, 1975; Wilches *et al*., 2016), but this could be a vulnerable life stage to target physical control methods such as sweeping or hand picking. Timely and accurate pest detection and identification is crucial for pest control. Automated artificial intelligence models are currently being tested for detection of crop pests, with detection accuracy of >90% (Fedor *et al*., 2009; Wang *et al*., 2023). Commercial pheromone monitoring lures are available for *D. maculatus*, and interspecific cross attraction of pheromone components is reported in some beetle species (Silva *et al*., 2018), however a species-specific lure could prove to be more attractive. *Dermestes ater* prefer to oviposit and pupate in crevices (Dr. M.J. Muzzatti, observations), and so this could be an important feature included in a trap design paired with pheromone monitoring. Biocontrol is an option, but again is difficult to achieve without adversely affecting the farmed insect. In museology, where dermestid beetles are reared for cleaning skeletal tissue, dermestids are extremely susceptible to mold, mites and other pests (Timm *et al*., 2020). The copra beetle (*Necrobia rufipes*) is a destructive predatory pest of dermestid colonies (Timm *et al*., 2020), but would be an unfavourable biocontrol agent for dermestids in mass reared insect production because it may also feed on the various life stages of the beneficial insect.

## Acknowledgements

We would like to thank Dr. Mads K Andersen for helping troubleshoot the hemocyte assays.

## Author contributions

MJM: Conceptualization; Data curation; Formal analysis; Funding acquisition; Investigation; Methodology; Project administration; Software; Validation; Visualization; Writing – original draft; Writing – review and editing. MWR: Conceptualization; Investigation; Methodology; Writing – review and editing. ECB: Methodology; Resources. SMB: Conceptualization; Funding acquisition; Resources; Supervision; Writing – review and editing. HAM: Conceptualization; Funding acquisition; Resources; Supervision; Writing – review and editing.

