## Supplemental tables and figures for "Farmed crickets raised with dermestids suffer from reduced and delayed growth, but not enough to explain reports of dramatic yield loss"

Colonel By Dr, Ottawa, Ontario, K16 5B6

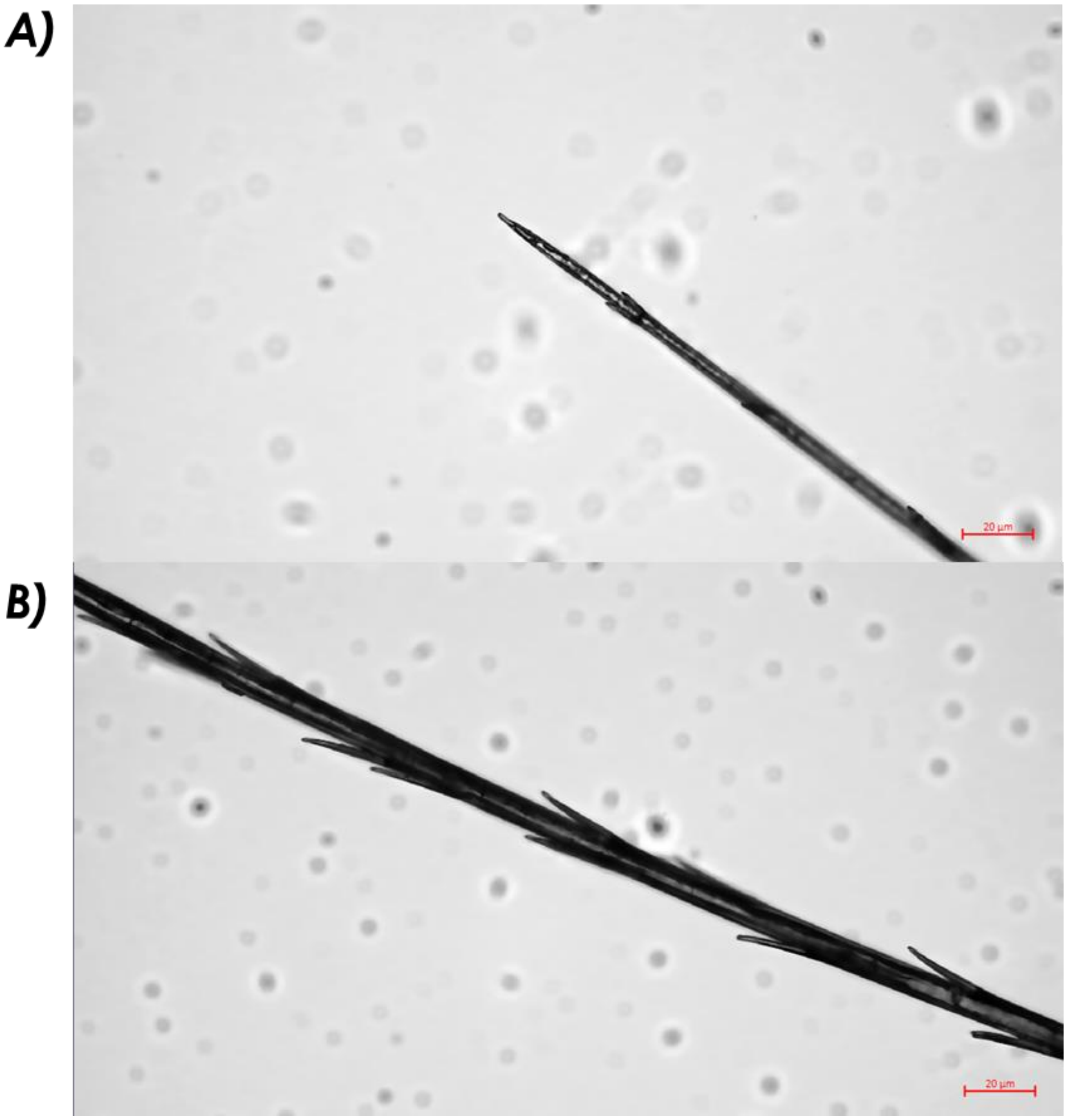


Figure S1. The tip (A) and the barbed sheath (B) of a hastisetae removed from a single late-instar *Dermestes ater* larva.

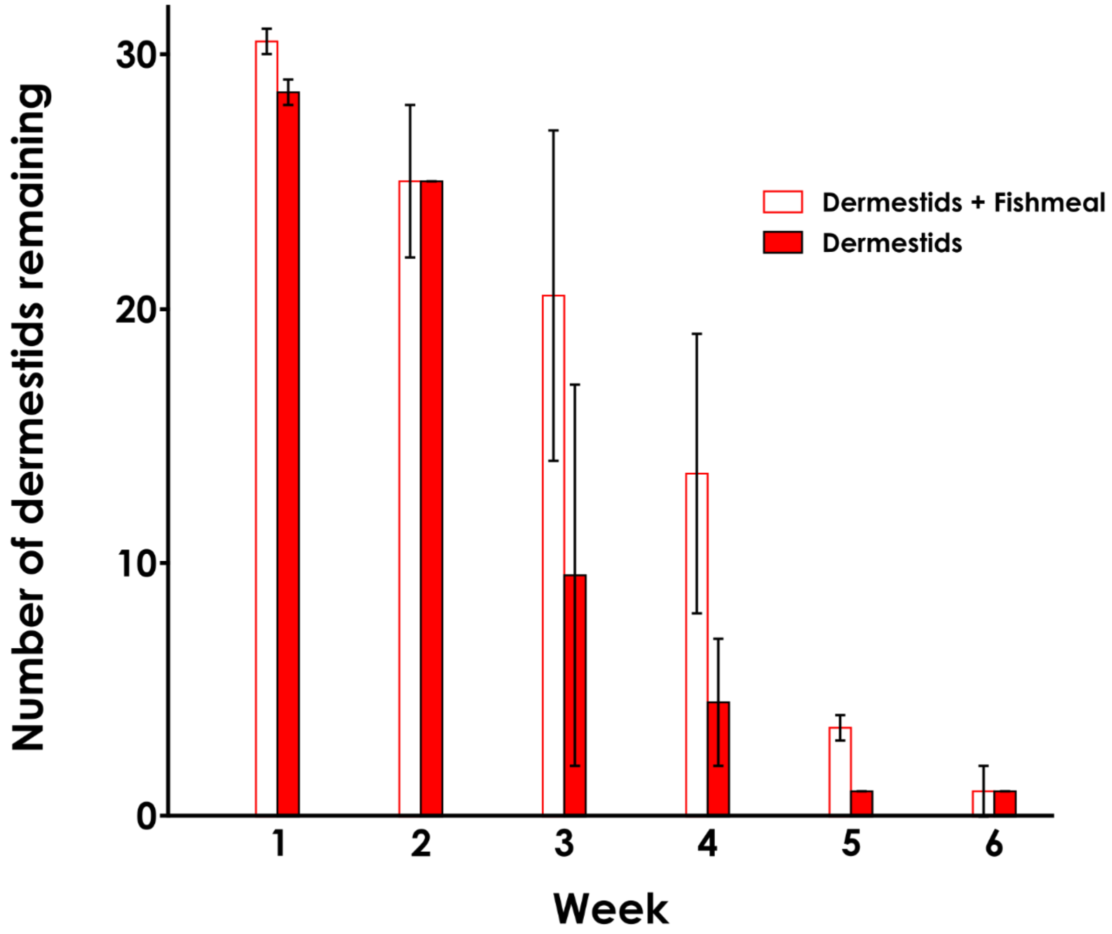


**Figure S2.** The number of dermestids remaining in each treatment from the competition experiment. Bins received N=30 late instar larvae every week, and at the end of each week all remaining dermestids (larvae, pupae, and adults) were counted and replaced. Error bars represent standard error.

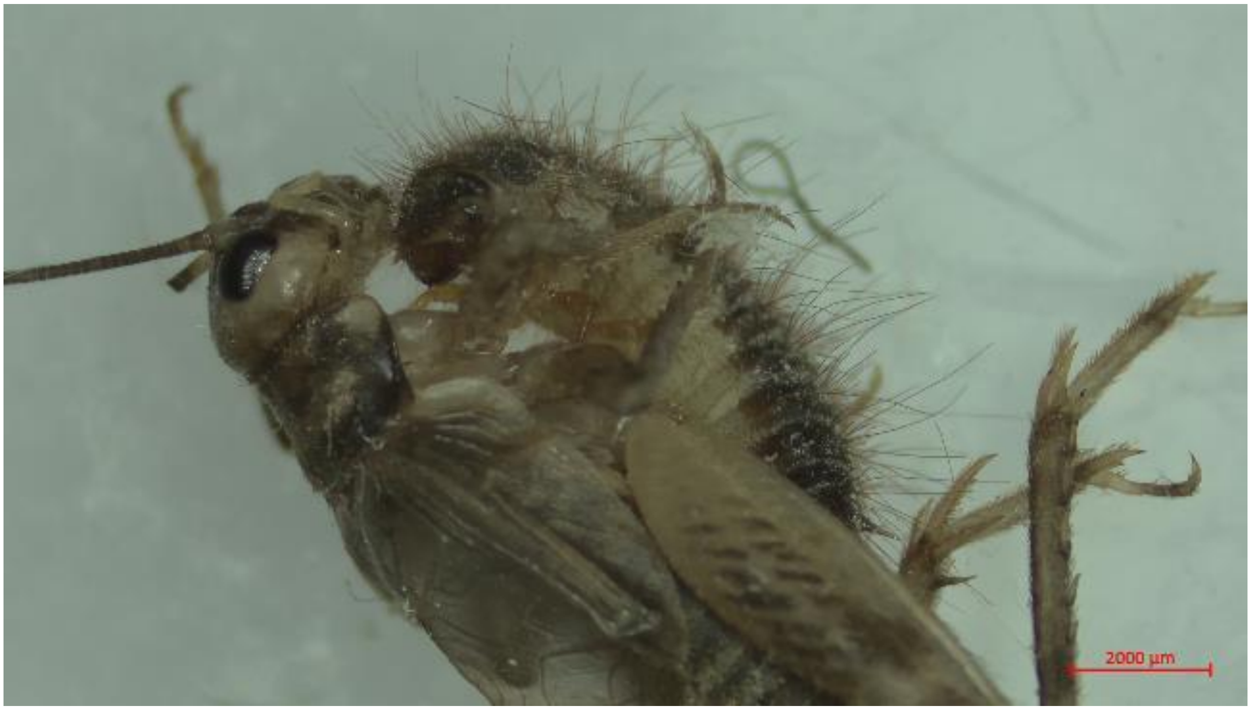


**Figure S3.** An adult male cricket (*Gryllodes sigillatus*) entangled by the hastisetae of a late instar *Dermestes ater* larva. The cricket was presumed dead based on unresponsiveness to tactile stimulation, and the dermestid larva was feeding on the cricket.

**Table S1.** Diet composition and relative protein and carbohydrate contributions of each ingredient for the competition experiment.

| | Conventional Cricket Diet | | | Fishmeal Removed Diet | | |
|---|---|---|---|---|---|---|
| Ingredient | As Fed (%) | Protein (%) | Carbohydrates (%) | As Fed (%) | Protein (%) | Carbohydrates (%) |
| Fish meal | 5.00 | 3.40 | 0.00 | 0.00 | 0.00 | 0.00 |
| Corn – dry ground | 42.00 | 2.90 | 33.43 | 41.00 | 2.83 | 32.64 |
| CFS soybean meal | 39.00 | 14.78 | 11.35 | 39.00 | 14.78 | 11.35 |
| Linseed meal | 5.00 | 1.60 | 2.40 | 5.00 | 1.60 | 2.40 |
| Nutri-mix Elite Grower DDG | 5.00 | 0.00 | 0.00 | 5.00 | 0.00 | 0.00 |
| Corn gluten | 4.00 | 2.40 | 0.60 | 10.00 | 6.00 | 1.50 |
| Total | 100.00 | 25.08 | 47.78 | 100.00 | 25.21 | 47.89 |

**Table S2.** Summary statistics from the linear model of food consumption of individual crickets fed hairy and hairless dermestids for seven days. Hairless dermestids represent dermestid larvae with their hastisetae removed, whereas hairy dermestids had their hastisetae unaltered. Terms in bold are statistically significant.

| Model | $R^2$ | Term | Estimate (± SE) | *t-value* | *P* |
|---|---|---|---|---|---|
| Dermestids consumed | 0.053 | **Intercept** | **4.72 ± 0.30** | **15.70** | **< 0.001** |
| | | **Hairy** | **-0.88 ± 0.43** | **-2.08** | **0.042** |

**Table S3.** Summary statistics from a linear model of circulating hemocyte concentration of crickets living with or without late-instar dermestid larvae and fed a diet with or without fishmeal.

| Model | $R^2$ | Term | *df* | *F* | *P* |
|---|---|---|---|---|---|
| Hemocyte concentration | 0.56 | **Treatment** | **1, 25.20** | **10.81** | **<0.001** |
| | | **Sex** | **4, 25.60** | **6.57** | **0.025** |
| | | Treatment*sex | 4, 25.22 | 1.78 | 0.16 |
